# A promptable foundation model enables automated multi-task dataset construction and real-time pose estimation in mice

**DOI:** 10.64898/2026.10.02.756254

**Authors:** Juan Cobos, Steeve Thirard, Marwen Belkaid, Jérémie Naudé

## Abstract

Behavioural experiments often hinge on moments when animals crowd together. In these situations, existing pose-estimation methods usually fail because detected crops include neighbouring animals, and their body parts are easily confused between individuals. We solved it with DeepLabSAM, a pipeline that segments and tracks every animal from a single text prompt using the SAM 3 foundation model, followed by a mask-gated pretrained pose estimation network from DeepLabCut. Across recordings of multiple interacting mice, the proposed method substantially reduces misassigned keypoints under heavy occlusion. Furthermore, we used DeepLabSAM to automatically construct the Multi-Task Mouse Behaviour (MTMB) dataset, comprising 14,300 frames spanning nine behavioural assays, with each instance annotated for bounding box, mask, identity, and pose. SuperAnimal pretrained pose models transferred unpredictably across this dataset. Conversely, MouseLite, our RF-DETR pose-estimation model trained on the MTMB dataset, matched their in-distribution accuracy, generalized better to unseen assays, and achieved real-time (faster-than-acquisition) inference. Combined, DeepLabSAM, MTMB dataset, and MouseLite provide a scalable framework for robust, cross-assay behavioural pose estimation of freely interacting mice.

## 1 Introduction

Behaviour is the highest-level output of the nervous system, and the resolution at which it can be measured bounds the questions that can be asked of it. For most of the history of the field that measurement was made by hand. An observer scored events from video, or, at best, a centroid was tracked through an arena. Both approaches are slow, coarse, and inconsistent between observers and laboratories.

In recent years, machine learning has lifted that bound. Markerless pose estimation places user-defined body parts on freely moving animals at human-level accuracy (1, 2). Multi-animal systems extend this to socially interacting animals (3, 4), and trackers follow individuals through long recordings without losing identity (5–7). Built on those measurements, unsupervised methods can now decompose continuous movement into recurring sub-second units without a human-defined ethogram (8, 9).

Yet these methods share a blind spot exactly where behaviour is most informative (10). When animals touch or overlap (i.e. the defining condition of social interaction), a detector’s crop of one animal contains parts of its neighbours, and a generic pose head places keypoints on the wrong body. Identity assignments leak between individuals at the precise moments an experiment is designed to capture (10). This is not a corner case to be patched but a central obstacle to measuring multi-animal behaviour, and it is compounded by a practical one: most pipelines still require training a dedicated model, with hundreds of hand-annotated frames, for every new species, behavioural assay, and lab condition (11).

Prior efforts such as SuperAnimal (12) have made important progress toward pretrained, plug-and-play pose estimation models. Yet the broader behavioural analysis ecosystem remains fragmented. Frameworks often use incompatible annotation schemas, keypoint definitions, coordinate conventions, and file formats, limiting the reuse of models and datasets across tasks and laboratories. To promote interoperability, we adopt the Common Objects in Context (COCO) standard (13), a widely used format for computer vision datasets, rather than introduce another custom representation.

Likewise, existing mouse behavioural datasets target individual vision tasks in isolation. SuperAnimal-TopViewMouse-5K focuses on pose estimation (1, 3, 12, 14–18); UDMT on identity tracking (19, 20); MouseSIS on instance segmentation (21); and CalMS21 on behaviour classification (22, 23). Each provides an important benchmark, yet none jointly annotates detection, segmentation, and pose on shared footage under a common standard.

This is not merely a limitation of existing datasets, but a reflection of how behavioural analysis has traditionally been approached. Locating animals, delineating their bodies, maintaining identity, and estimating pose are mutually informative tasks, yet neuroscience workflows typically solve them independently using specialized software. Modern computer vision instead treats detection, segmentation, and keypoint estimation as a unified prediction problem under shared annotation standards and model architectures (13, 24, 25). We adopt that perspective here.

The remaining obstacle is not composing these tasks but doing so reliably when animals interact and become visually ambiguous. Rather than considering segmentation and keypoint tracking as distinct tasks for extracting behaviour from video as classically considered (10), here we hypothesized that accurate instance segmentation provides a natural intermediate representation. By explicitly separating overlapping animals, segmentation resolves the ambiguities that propagate to identity tracking and pose estimation. For example, promptable segmentation foundation models of the SAM family (26–28) have been adapted to medical imaging (MedSAM (29)) and microscopy (*µ*SAM (30)). We sought to take advantage of SAM 3 (28), which detects and tracks animals from a text prompt alone, and chain its segmentation to pose estimation: each animal’s instance mask gates the input of a pretrained pose estimator, so that the estimator sees only the animal whose keypoints it predicts.

Hence, we introduce DeepLabSAM, a unified behavioural analysis pipeline for detection, instance segmentation, and pose estimation. Using this pipeline, we assemble a multi-task behavioural dataset spanning nine widely used mouse assays that jointly annotates detection, instance segmentation, and pose under a common compatible format. We further train a family of compact RF-DETR (31) models, termed MouseLite, on this dataset. As a result, MouseLite retains much of DeepLabSAM’s robustness while dramatically reducing computational cost. Together these resources provide an accurate, standardized, and ready-to-use framework for large-scale behavioural analysis.

## 2 Results

### 2.1 Mask-gating prevents keypoint cross-assignment under occlusion

Measuring the behaviour of interacting animals requires keypoints that stay attached to the right individual, frame by frame (10). The difficulty is not detecting animals or placing keypoints on an isolated one (pretrained models do both well) but keeping the two joined when animals interact. A pipeline that solves this without manual annotation (which goes from hundreds to thousands of frames for every new species, camera view, and task) would remove the main obstacle to building behavioural datasets at scale.

Our solution, DeepLabSAM (Figure 1), streams a video through SAM 3 (28) which, from a single text prompt (e.g. “mouse”), detects every animal, segments it, and maintains its identity across frames as a persistent track. Each tracked instance is then cropped and passed to a pose head (the SuperAnimal top-view mouse model (12)). Because SAM 3 already assigns a track identity to each mask, each set of keypoints inherits that identity, with no need for a post-hoc step to match poses back to tracks.

**Figure 1:**
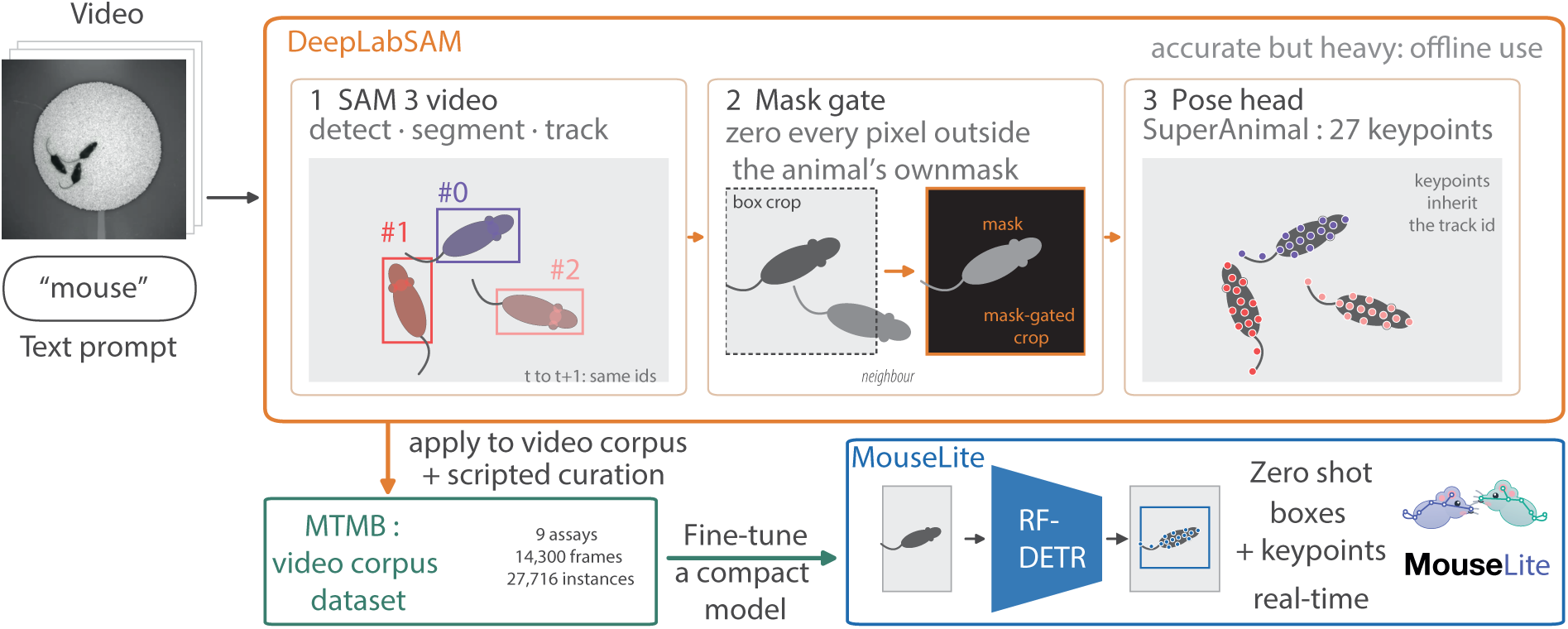
DeepLabSAM converts raw multi-animal video into identity-resolved keypoints from a single text prompt. The top band shows the three stages of DeepLabSAM; the bottom band shows its use as an annotation engine: applied to a video corpus with scripted curation, it produces the MTMB dataset, which is distilled into MouseLite, a single-stage real-time model. A raw video and a text prompt (“mouse”) are the only inputs. **SAM 3 Video** (28) detects every animal matching the prompt, returns a pixel-accurate instance mask and box for each, and propagates a persistent track identity across frames (coloured overlays; colour denotes track identity). **Mask gate** then isolates each tracked instance: its box is cropped into a 256 *→* 256 letterbox and every pixel outside the instance’s own mask is zeroed, so each crop contains one animal against a blank background. The **pose head** (SuperAnimal top-view mouse (12)) regresses 27 keypoints per crop. The output is a per-frame set of boxes, masks and poses, each carrying the track identity assigned by SAM 3, so keypoints are identity-resolved without any post-hoc assembly step.

Two problems separate the naive composition from a usable tool. First, a pose head run on a bounding box-crop has no notion of identity, so when a neighbour intrudes into the crop, the pose estimator places keypoints on the wrong animal (Figure 2). Such failures become frequent in crowded or complex scenes (Figure 2c,d). Second, the composition presents a scaling problem: SAM 3 retains a memory of every tracked frame, so on long recordings GPU memory grows without bound until inference fails. We resolve both issues with **mask-gating** (Figure 2a, b) and with an additional **bounded mask-memory** fix, which keeps GPU memory flat on long videos at no measurable runtime cost (Figure S1).

**Figure 2:**
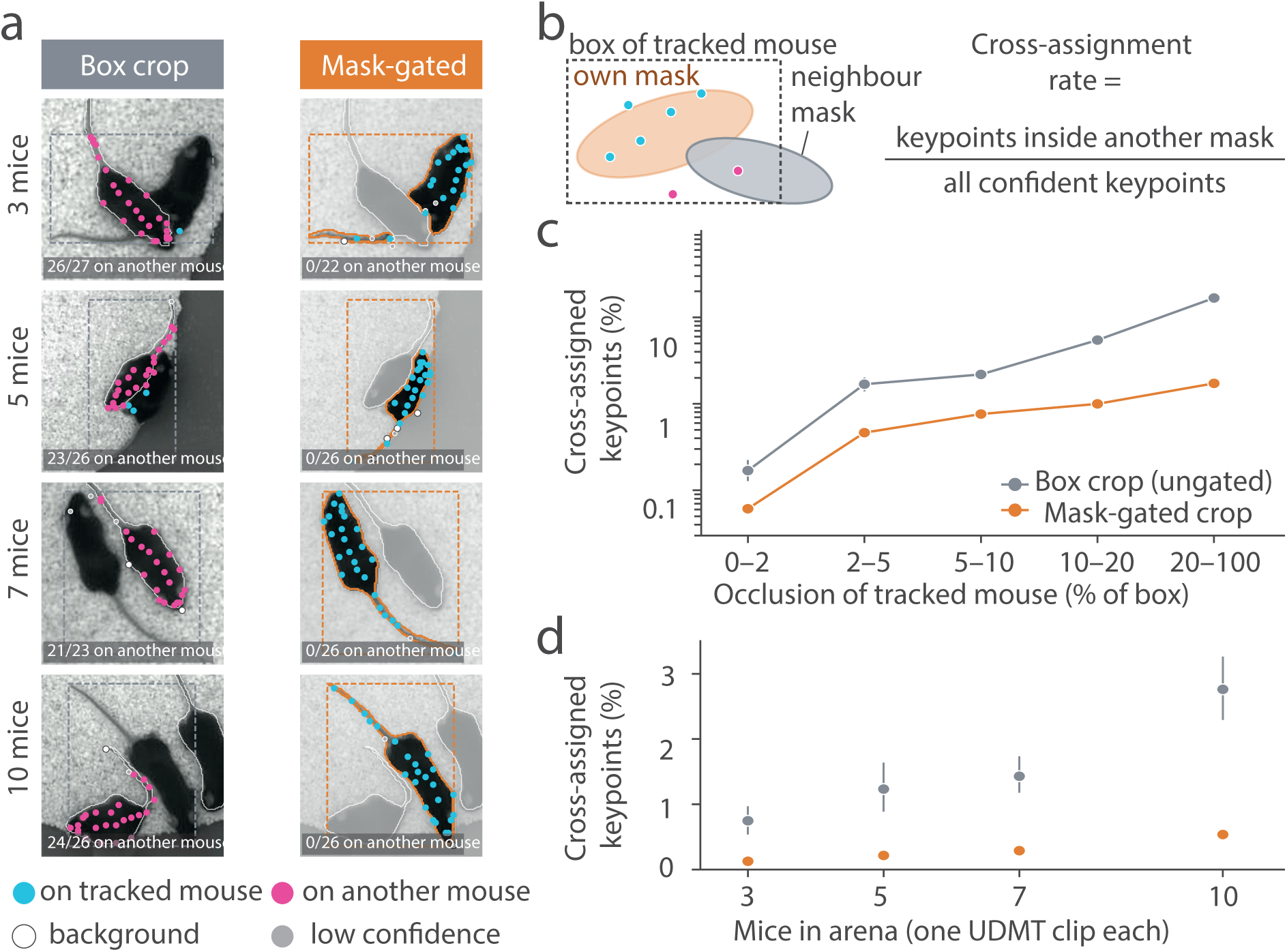
Mask-gating prevents keypoint cross-assignment under occlusion. All panels use the four public UDMT mouse clips of 3, 5, 7 and 10 freely interacting animals (19, 20). Throughout, grey is the baseline (ungated box crop) and orange the proposed mechanism (mask-gated crop). (**a**), Selected examples, one per clip, illustrating the failure mode. Left, box crop: keypoints are placed on a neighbouring mouse (magenta). Right, mask-gated crop, in which pixels outside the tracked mouse’s mask are suppressed (faded): keypoints stay on the tracked mouse (cyan). Dashed box, tracked box. Orange outline, tracked mask. White outlines, neighbours’ masks. Inset text, confident keypoints falling inside another mouse’s mask. (**b**), Cross-assignment rate: the percentage of confident keypoints (confidence > 0.3) that fall inside another animal’s mask. These keypoints are wrong by construction, so no ground truth is needed. Keypoints on animals without a mask count as background, making the rate a lower bound. (**c**), Cross-assignment rate as a function of the occlusion of the tracked mouse (fraction of its box covered by other masks; right-open bins), all clips pooled. Log scale. (**d**), The same measure per clip (151,963 tracked instances in total). In c and d, rates are pooled within each stratum (summed cross-assigned keypoints over summed confident keypoints). Error bars are 95% confidence intervals from a paired bootstrap (2,000 resamples of 10-s blocks of frames within each clip); where not visible, they are smaller than the marker.

We characterized both mechanisms on four publicly available recordings from the UDMT (Unsupervised Deep-learning for Multi-animal Tracking) dataset (19, 20), respectively involving 3, 5, 7 and 10 mice interacting freely in the same arena. Because the clips differ only in how many animals they contain and are several thousand frames long, we can vary the two quantities the mechanisms act on — how crowded the scene is, and how long the video runs — while holding everything else fixed.

We characterized both mechanisms on four publicly available recordings from the UDMT (Unsupervised Deep-learning for Multi-animal Tracking) dataset (19, 20), respectively involving 3, 5, 7 and 10 mice interacting freely in the same arena. Because the clips differ only in how many animals they contain and are several thousand frames long, we can vary the two quantities the mechanisms act on — how crowded the scene is, and how long the video runs — while holding everything else fixed.

To isolate the mask gate’s effect, we ran the pose head twice on the same SAM 3 boxes, once with the mask applied and once without, so that the only variable is whether neighbouring animals are zeroed out. Without keypoint ground truth on these clips, we measured **cross-assignment rate**: the fraction of an animal’s confident keypoints that land inside a *different* animal’s mask, which are wrong whatever the true pose. As expected, where animals are well separated the mask-gating changes almost nothing (0.17% against 0.06% cross-assignment below 2% occlusion). Its effect grows with crowding and occlusion, cutting cross-assignment from 16.7% to 1.7% above 20% in the most occluded frames (Figure 2c). The same reduction holds across every clip from three to ten mice (Figure 2d). Mask-gating thus removes most of the errors introduced by crowding. Cross-assignment is computed frame by frame, so it measures whether keypoints land on the correct animal at each instant; it does not assess whether a track follows the same animal across frames, which depends on SAM 3’s mask propagation.

SAM 3’s default streaming state keeps a memory entry for every tracked frame and object (28, 32), and out-of-memory failures have been reported for multi-animal sequences even on a 96-GB GPU (33). We cap SAM 3’s mask-memory to a fixed span and evict older entries (Methods). Peak GPU memory then stays flat regardless of video length, whereas the default grows linearly with both duration and animal count (Figure S1a,b), and the cap adds no measurable runtime overhead (Figure S1c). Throughput is instead set by SAM 3 itself. Considering only its detection and tracking, before the pose head, it runs at a few frames per second and slows further as animals accumulate (Figure S1d). Finally, DeepLabSAM’s keypoints were compared with human annotations on the test split of SuperAnimal-TopViewMouse-5K (12) (Figure S2). Because the SuperAnimal pose head was trained on this dataset, this comparison is in-distribution for the pose head: it tests the SAM 3 and mask-gating front end, not the transfer of the pose head to new footage.

DeepLabSAM is therefore best used as an *o“ine* annotation engine: run once over a corpus of behavioural video, it produces identity-resolved, occlusion-robust keypoint labels without any manual annotation, the regime in which throughput is amortized and the elimination of human labelling is the decisive advantage.

### 2.2 A multi-task dataset built by DeepLabSAM

Until now, the cost of labelling crowded videos has made it difficult to assemble unified datasets, leaving existing resources fragmented across species, viewpoints, and labelling conventions (3, 4, 12, 19, 23, 34). We therefore used DeepLabSAM to create a dataset no existing resource provides: detection, segmentation, tracking, and pose annotations on the same frames.

The Multi-Task Mouse Behaviour (MTMB) dataset comprises 14,300 frames and 27,716 annotated mouse instances at 1280 *→* 720, drawn from nine standard behavioural assays (Table 1, Figure 3a): Barnes maze, direct interaction, marble burying, novel-object recognition (two variants), open field, social interaction, T-maze, and three-chamber sociability. Every instance carries a bounding box, a pixel-accurate instance mask, a persistent track identity, and a 27-point pose in the SuperAnimal top-view mouse convention (12), all unified under the COCO format (13). To our knowledge, it is the first behavioural dataset to unify these annotations on real assay footage, rather than distributing the tasks across disjoint benchmarks.

**Figure 3:**
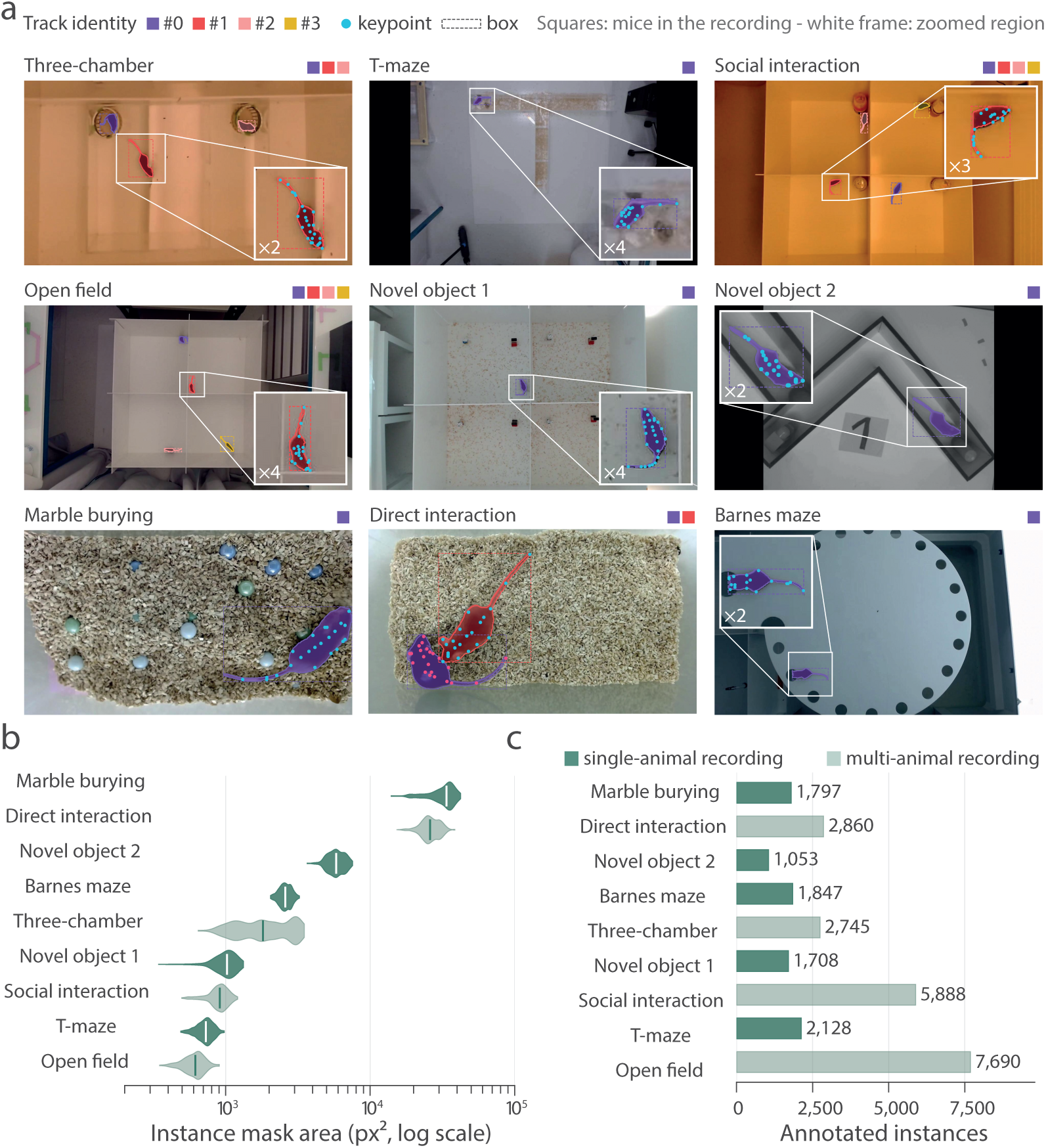
The Multi-Task Mouse Behaviour (MTMB) dataset spans nine assays. (**a**), Representative frames from the nine recordings, re-rendered from the curated annotations. Each instance carries a segmentation mask (filled), a bounding box (dashed) and 27 keypoints (cyan; visible keypoints only). Colour encodes the tracked identity (mouse #0–#3) and is consistent within a recording. Squares above each frame give the number of mice in the recording. Where animals are small, keypoints are shown in the magnified inset (white frame; magnification indicated), which corresponds to the region outlined in the full frame. (**b**), Distribution of instance-mask area per recording (log scale), ordered by median (vertical tick). Median area ranges from 617 px² (open field, four mice) to 33,763 px² (marble burying), a 55-fold span. Dark, single-animal recordings; light, multi-animal recordings. (**c**), Number of annotated instances per recording (27,716 in total), with the number of mice per recording at right. The four multi-animal recordings contribute 19,183 instances (69%).

**Table 1:** Dataset composition. One row per task, each task a single continuous recording labelled frame by frame. *Frames* counts the labelled frames, *Instances* the animal instances annotated in them, and *Animals* the mice a recording holds, read off the frame carrying the most instances. *Duration* is the frame count at 30 fps and *Median box* the median area of an annotation’s bounding box.

| Task | Frames | Instances | Animals | Duration | Median box (px<sup>2</sup>) |
| --- | --- | --- | --- | --- | --- |
| Barnes maze | 1,847 | 1,847 | 1 | 62 s | 8,241 |
| Direct interaction | 1,430 | 2,860 | 2 | 48 s | 92,987 |
| Marble burying | 1,797 | 1,797 | 1 | 60 s | 102,820 |
| Novel object 1 | 1,719 | 1,708 | 1 | 57 s | 2,992 |
| Novel object 2 | 1,053 | 1,053 | 1 | 35 s | 22,518 |
| Open field | 1,939 | 7,690 | 4 | 65 s | 1,377 |
| Social interaction | 1,472 | 5,888 | 4 | 49 s | 2,352 |
| T-maze | 2,128 | 2,128 | 1 | 71 s | 1,409 |
| Three-chamber | 915 | 2,745 | 3 | 30 s | 2,816 |
| **Total** | **14,300** | **27,716** |  | **7.9 min** |  |

Every label in the dataset was generated automatically by DeepLabSAM over each recording, followed by human curation. Curation is scripted and auditable rather than frame-by-frame retouching — masks repaired by connected components, re-acquired identities merged, lead-in frames dropped (Figure S3) — so the labels are reproducible from the raw video and the corrections themselves are recorded (Methods).

The nine recordings span a wide range of conditions that would prevent direct, trivial transfer. Apparent animal size varies 55-fold across assays, from a median instance-mask area of 617 px^2^ in the four-animal open field to 33,763 px^2^ in marble burying (Figure 3b). Animal counts run from one to four, and arenas, lighting, substrate, and occlusion vary from bare open-field to deep bedding (Figure 3a). Four of the nine recordings are multi-animal, contributing 19,183 instances (69% of the dataset) recorded under crowding, the regime where identity-preserving annotation matters most and where naive detect-then-crop pipelines fail. We then leveraged this variety in downstream models that can learn robustness from it.

### 2.3 Zero-shot pose transfer varies widely across real behavioural assays

A dataset spanning real assays lets us ask how far do pretrained models carry across the range of conditions a behavioural lab meets. We evaluated the standard pretrained SuperAnimal top-view mouse (12) on every tenth frame of the nine recordings (1,433 frames, 2,778 instances) and scored against the curated labels with Object Keypoint Similarity (OKS) mean average precision (13).

This evaluation is deliberately zero-shot: no fine-tuning and no video adaptation, both of which DeepLabCut recommends when deploying to a new domain. The reported performance therefore represents an out-of-the-box baseline rather than the model’s achievable ceiling.

Throughout this evaluation the pose estimation network is held fixed. The reference labels are generated by DeepLabSAM, then curated. Scores on MTMB therefore measure agreement with these curated labels; DeepLabSAM’s agreement with human annotations was measured on the SuperAnimal-TopViewMouse-5K test split (OKS AP 0.696, median OKS 0.965; Figure S2), a benchmark that is in-distribution for the pose head (Section 2.1). The predictions being scored come from the same pose network but in the pretrained model’s native form, DeepLabCut’s Faster R-CNN detector on rectangular crops (12, 35). Because labels and predictions therefore share an identical pose head and differ only in detection and cropping, this has two consequences. First, any bias the pose head carries is shared by predictions and annotations and cancels in OKS, so the reported values are a conservative lower bound on what a better front end can recover. Second, the differences observed below should be attributed to detection and cropping rather than to the pose estimator.

Zero-shot performance nevertheless varied markedly across assays (Figure 4a,b). Keypoint mAP reached 0.86 on the Barnes maze, 0.68 and 0.59 on the two novel object recognition sessions, and 0.47 on direct interaction, but fell to 0.23 in the open field and below 0.10 on the three-chamber social test (0.09), marble burying (0.07) and the T-maze (0.02). On the social interaction recording the detector returned no instances in any of its frames, giving 0.00. The pretrained model therefore generalized well to several assay types without adaptation while failing outright on others, spanning complete failure to near-ceiling accuracy within a single benchmark.

**Figure 4:**
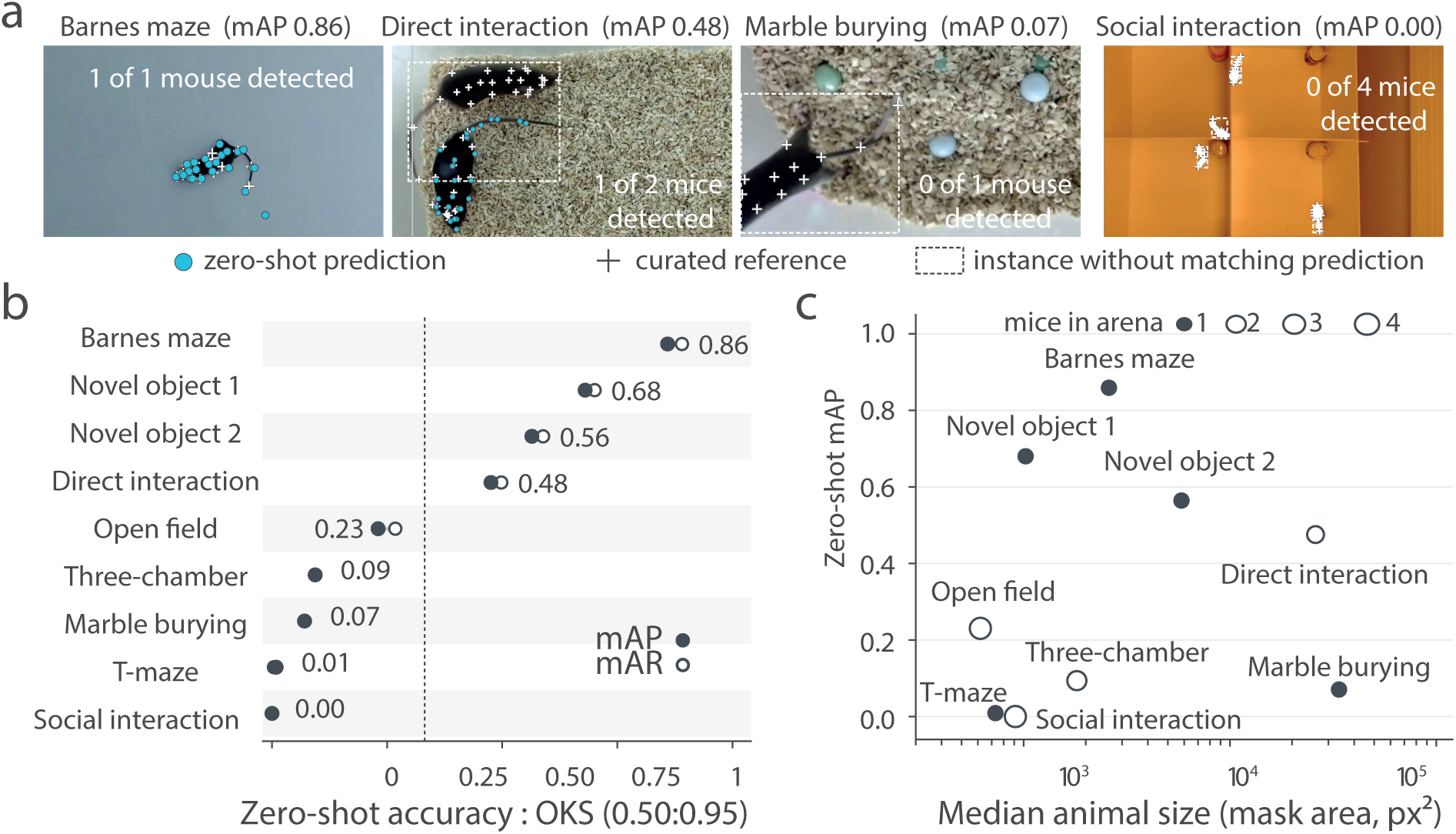
Zero-shot pretrained SuperAnimal transfers unevenly across standard behavioural assays. We evaluated the pretrained model (12) in its native form across nine recordings, scoring predictions against curated labels using OKS with a uniform keypoint sigma of 0.1. Recordings are ordered by mAP in both panels. (**a**), Zero-shot predictions (cyan dots) against the curated reference (white crosses) on the first evaluated frame of four assays. Dashed boxes mark reference instances without a matching prediction. Inset text gives the detected instances and, for single-animal frames, the OKS of the prediction. (**b**), Keypoint mAP (filled) and mAR (open) per assay, OKS 0.50:0.95. Dashed line, mean mAP over the nine assays. In social interaction the detector returned no instance in any frame. (**c**), Zero-shot mAP against median animal size (instance-mask area, log scale). Marker size gives the number of mice in the arena; filled markers are single-animal recordings. Neither size nor animal count predicts transfer.

The strongest-performing recordings also maintained high accuracy under stricter localization criteria (Figure 4b). On the Barnes maze, AP_50_ and AP_75_ both reached 1.00, and the two novel-object variants achieved 0.75 and 0.71 at AP_50_ and 0.74 and 0.71 at AP_75_. Direct interaction (0.67 and 0.55) and the open field (0.32 and 0.25) lost more accuracy when the threshold was tightened, yet still retained roughly 80% of their AP_50_ (Figure S4a). These results indicate that successful zero-shot transfer is not merely due to coarse localization: when the detector isolates the animal, the pretrained pose estimator also produces anatomically consistent keypoints.

In contrast, three-chamber social testing (AP_50_ = 0.12), marble burying (0.10) and especially the T-maze (0.01) remained poor even at the most permissive threshold, and the social interaction recording yielded no correct instances at all. Average recall closely tracked average precision in every assay (Figure S4b), so the detections that were returned were almost always correct and the losses came from missed animals. Because the poor assays fail at AP_50_ and not only at AP_75_, the dominant failure is recovering suitable animal instances rather than refining keypoint geometry.

No single recording characteristic predicts where an assay falls within this range. Poor performance occurred for both large animals (marble burying) and relatively small ones (T-maze), while the smallest animals in the benchmark (the four-animal open-field recording) achieved intermediate performance. Likewise, both single-animal (T-maze, marble burying) and multi-animal (direct interaction, open field) recordings appear throughout the ranking. Assay identity alone therefore provides little indication of whether a pretrained model will transfer successfully. Instead, zero-shot performance appears to depend on how closely the recording conditions resemble those represented during pretraining.

Fine-tuning, video adaptation, and improved detection front ends are all expected to reduce this variability, but each requires representative annotations from the target assay. The dataset introduced above provides precisely these annotations, enabling the adaptation experiments presented in the following section.

### 2.4 A real-time single-stage pose model matches state-of-the-art accuracy and generalizes better to unseen assays

While the adapted foundational model DeepLabSAM achieves strong zero-shot accuracy without labels, it does so at the cost of high latency, and current models for behavioural analysis generalize unpredictably to new real-world assays. Filling this gap requires a general tool with transfer capabilities across different recording conditions and setups while maintaining sufficient processing speed and portability.

Thus, we trained RF-DETR (31), a single-stage transformer architecture (36) that predicts boxes and keypoints jointly, on our Multi-Task Mouse Behaviour dataset. Then, we compared the resulting model, termed MouseLite, against the DeepLabCut cascade (Faster R-CNN detector followed by the SuperAnimal top-view pose head (12, 35)) fine-tuned on the same dataset.

To test each model’s ceiling performance, both were trained and evaluated on frames drawn from all nine recordings, measuring in-distribution accuracy. In this condition, the two fine-tuned models are close, with the advantage splitting by task (Figure 5a). On detection, MouseLite leads, reaching 0.85 box mAP against the cascade’s 0.72, a 13-point margin that reflects the single-stage detector’s stronger localization on this footage. On keypoints the ordering reverses but the gap narrows: the cascade scores 0.88 OKS mAP and MouseLite 0.83, the dedicated pose head retaining a small edge on the task it specializes in. In-distribution, then, the single-stage model is competitive with the fine-tuned cascade on both tasks and superior on detection.

**Figure 5:**
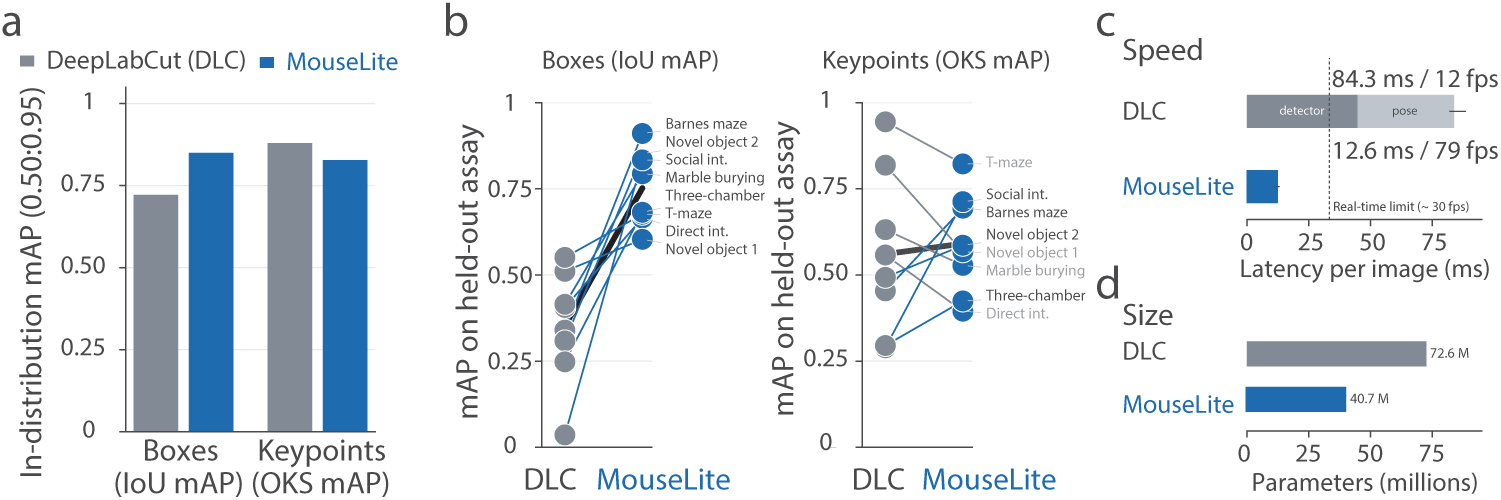
MouseLite, trained on MTMB, doubles detection accuracy on unseen assays and matches keypoint accuracy, being markedly cheaper and faster to run. (**a**), In-distribution accuracy (pooled split): box mAP (IoU 0.50:0.95) and keypoint mAP (OKS 0.50:0.95). (**b**), Generalization to unseen assays (leave-one-assay-out, n = 8 held-out assays). Each line joins DeepLabCut (DLC) and MouseLite on the same held-out assay. Blue lines mark assays where MouseLite scores higher; grey lines mark the reverse. The shaded line joins the means. MouseLite achieved higher box mAP on all eight assays (mean 0.35 → 0.75; two-sided Wilcoxon signed-rank test, P = 0.008). Keypoint mAP was comparable (0.56 → 0.59; MouseLite higher on 4/8 assays; P = 0.95). (**c**), Inference latency per image: median (bar) and 5th–95th percentile across images (whisker). Measured on 185 test images of the Barnes maze fold, batch size 1, fp16, after 30 warm-up iterations, on a single NVIDIA RTX 4000 Ada GPU [to be confirmed for MouseLite]. MouseLite needs 12.6 ms (79 fps). The two-stage cascade needs 84.3 ms (12 fps), split into detector (44.8 ms) and pose estimation (39.0 ms). Dashed line, 30 fps video rate. (**d**), Model size: 40.7 M parameters for MouseLite versus 72.6 M for the cascade.

Next, to assess transfer capabilities, we used a *leave-one-assay-out* protocol in which each assay is held out entirely for testing, a second assay is held out for validation, and the models train on the remaining seven (Methods). Because each assay is represented by a single recording, this protocol measures transfer to an unseen recording of an unseen assay type. The open-field fold is not included in the comparison because of a software failure of the DeepLabCut cascade on this fold. Here MouseLite showed higher accuracy (Figure 5b). Averaged over the eight held-out assays, MouseLite reaches 0.75 box mAP against the cascade’s 0.35, more than doubling detection accuracy on unseen assays, and 0.59 keypoint mAP against 0.56. The per-assay scatter (points in Figure 5b) shows the gap is systematic rather than driven by a few folds. On keypoints the two are closer on average and each wins on some folds, but MouseLite’s spread is narrower and its worst cases less severe. Even when both models are fine-tuned on the same training assays, the single-stage detector generalizes to held-out assays far better than the cascade.

Beyond accuracy, the single-stage design is markedly cheaper to run. On a single GPU, MouseLite processes an image in 12.6 ms (79 fps) against the cascade’s 84.3 ms (12 fps), a 6.7*→* speed-up that clears the threshold for real-time analysis (Figure 5c), and it does so with fewer parameters: 40.7 M against the cascade’s 72.6 M (Figure 5d). The two designs also scale differently with group size: MouseLite predicts boxes and keypoints for all animals in a single forward pass, whereas the cascade runs its pose head once per detected animal, so its cost grows with the number of animals in the frame, a known limitation for live multi-animal tracking (10).

In conclusion, the single-stage model matches the cascade in-distribution, surpasses it in detection on unseen assays while matching its keypoint accuracy, and runs substantially faster on a smaller footprint — the combination of accuracy, generalization, and speed needed for a portable, general-purpose tool. Alongside this joint detection–pose model, we also release MouseLite–detection and MouseLite–segmentation models trained on the pooled dataset across a range of scales, to cover deployment settings with different accuracy–latency trade-offs (Figure S5).

## 3 Discussion

We asked whether the foundation-model strategy that has transformed adjacent domains could be adapted to behavioural video, where the goal is not simply to recognize images but to quantify the continuous and socially complex dynamics of living organisms. Rather than train task-specific models from scratch, we adapted promptable segmentation foundation models from the SAM family (26–28), following the successful development of domain-adapted foundation models in medical imaging (29) and microscopy (30). Our results show both the promise and the limits of this approach: foundation models can provide a powerful annotation engine for behavioural analysis (10), but practical deployment requires additional specialization.

Behavioural video differs from the image domains in which foundation models have flourished in two fundamental ways. First, the unit of analysis is not an image but a continuous recording: minutes to hours of video make memory footprint and throughput central constraints rather than secondary engineering details. Second, behaviour cannot be reduced to segmentation alone. Meaningful analysis requires identities maintained across long interactions (5–7) and anatomical landmarks that describe movement (1, 3, 4). DeepLabSAM addresses these requirements by combining SAM 3 (28, 32) for promptable video segmentation with a mask-gated pose head derived from the DeepLabCut SuperAnimal models (12).

A key advantage of this design is modularity (Figure 1). The segmentation component defines individual instances and can, in principle, route different pose models to different tracked objects, enabling analysis across diverse organisms and experimental settings without redesigning the entire pipeline. However, quantitative evaluation in new organisms requires curated references and remains an important direction for future work.

At the same time, our results highlight a practical asymmetry of current foundation models. They can be sufficiently accurate for annotation while remaining too computationally demanding for routine deployment. DeepLabSAM’s resource requirements, restrictive licensing, and limited throughput at high resolution make it unsuitable for closed-loop experiments (37) or everyday analysis of large behavioural datasets. This motivates our strategy of using DeepLabSAM as an annotation engine and training lightweight models for practical deployment.

The Multi-Task Mouse Behaviour dataset (Figure 3) embodies this strategy by placing detection, instance segmentation, tracking, and pose annotations on the same recordings under a unified schema. Existing behavioural resources provide valuable benchmarks for individual tasks (19, 23, 34, 38, 39), but rarely allow these complementary views of the same animal to be studied jointly. The resulting dataset can therefore support both task-specific evaluation and multi-task model development across diverse assays, imaging conditions, animal sizes, and social contexts.

The dataset also exposes an important limitation of current pretrained models: strong performance on one behavioural domain does not guarantee transfer to new assays or experimental conditions. Zero-shot evaluation revealed substantial degradation outside the conditions represented during training, whereas fine-tuning recovered performance and established a realistic baseline using SuperAnimal models (12). This motivated the development of compact RF-DETR-based models (31), which retain much of the accuracy of larger systems while substantially improving throughput and accessibility. In particular, MouseLite generalized to held-out assays, suggesting that carefully constructed multi-task datasets can reduce the need for experiment-specific annotation.

Together, these results establish a general framework for behavioural analysis in which foundation models provide scalable annotation, and lightweight specialized models enable efficient deployment. Extending this approach beyond mice will require additional data across species, but the shared annotation standard and modular design create a foundation for building interoperable behavioural models at scale.

## 4 Methods

### 4.1 Animals

Male C57BL/6J mice, 10 weeks of age, were used in this study. Animals were housed in groups of 2–5 mice per cage under standard laboratory conditions, with a 12-h light/dark cycle, *ad libitum* access to food and water, a controlled temperature of 22 *±* 2 *^→^*C, and relative humidity of 55 *±* 10%. All animal procedures were conducted in accordance with the guidelines of the French Ministry of Agriculture and Forestry for the care and use of laboratory animals and were performed under the authorized establishment license B34-172-41. Prior to behavioural testing, mice were habituated to the experimenter for 7 days. Before each behavioural test, animals were allowed to habituate to the experimental room for 30 min.

### 4.2 Public datasets

#### 4.2.1 UDMT

Mask-gating and mask-memory eviction were characterized on four clips of the public UDMT mouse recordings (19, 20), each showing 3, 5, 7, or 10 freely interacting mice in a circular arena filmed from above. The clips are 600–664 px square, 6–8 min long, and were resampled to 15 fps. Every reported measure is either label-free (cross-assignment) or a hardware measurement (memory, runtime), so none requires ground-truth annotation.

#### 4.2.2 SuperAnimal-TopViewMouse-5K

SuperAnimal-TopViewMouse-5K (12) aggregates keypoint annotations from a dozen source datasets collected by multiple laboratories (1, 3, 14–18). The test split from this dataset was used to evaluate the zero-shot predictions generated by DeepLabSAM against human-annotated ground truth (Figure S2). Because the SuperAnimal pose head used in DeepLabSAM was trained on this dataset, this evaluation is in-distribution for the pose head and mainly tests the SAM 3 and mask-gating front end.

### 4.3 DeepLabSAM architecture

DeepLabSAM streams a video through SAM 3 (28) and passes each tracked instance to a pose head; no component is trained.

#### Detection, instance segmentation and tracking

Frames are decoded on the GPU and fed one at a time to the SAM 3 video model (`facebook/sam3`, bfloat16, 1008 *→* 1008 input) with a single text prompt (“`mouse`“). Each frame returns one mask, box, detection score and persistent track identity per matching instance. The inference session is reset per video, so prompts and tracking state never leak between recordings.

#### Bounded mask-memory

SAM 3’s streaming state keeps the mask-memory of every previously tracked frame for every object. After each forward pass we delete, per object, all non-conditioning memory entries older than the current frame index minus a fixed window (e.g. 64 frames), together with any reconditioned conditioning frames older than the same cut-off. The first conditioning frame is always retained as the anchor the model requires. Setting the window to None restores SAM 3’s default unbounded behaviour and is the *eviction off* condition throughout. A comparable sliding-window memory has been used for SAM 2-based multi-object tracking (40).

#### Mask-gated pose

Pose is estimated with the DeepLabCut SuperAnimal `topview-mouse` model (12) (HRNet-W32 backbone, 27 keypoints), run on the SAM 3 boxes rather than on its own detector. Each box is cropped from the frame, resized with aspect preserved and centre-padded to a 256 *→* 256 letterbox, then normalized with ImageNet statistics. The instance’s own mask is letterboxed identically and multiplied into the crop after normalization, so the pose head sees only the intended animal, not overlapping neighbours or background. Keypoints are decoded from the heatmap head and mapped back to image coordinates by inverting the letterbox. Both mask-gated and ungated conditions therefore share boxes, crops and weights, and differ only in the gating mechanism.

### 4.4 Multi-Task Mouse Behaviour Dataset

Nine continuous recordings of standard behavioural assays (Barnes maze, direct interaction, marble burying, two variants of novel-object recognition, open field, social interaction, T-maze and three-chamber sociability) were filmed from above and decoded to consecutive 1280 *→* 720 frames at 30 fps. In total there are 14,300 frames corresponding to *↑*7.9 min of footage (see Table 1).

Annotations are model generated and split into two passes over each recording — instance segmentation and tracking first, pose second — with a curation step after each, so that identity and mask errors are corrected before they can propagate into the poses that are conditioned on them.

#### Pass 1: detection, instance segmentation and tracking

The recording is streamed frame by frame through the SAM 3 video wrapper with the prompt “`mouse`“ at 1008 px. Every frame is passed through the tracker to preserve continuity. Each frame is written as a JPEG together with a COCO entry per instance carrying a box, an exact instance segmentation (compressed RLE for masks with holes or several parts, polygon otherwise), the mask pixel count as area, SAM 3’s detection score, and the persistent `track_id`. The *K* highest-scoring instances per frame are kept matching the number of animals in the arena. Per-frame overlays were rendered alongside for visual inspection.

#### Curation 1: masks, identities and frames

Before a recording joins the dataset, its instance segmentation file is repaired programmatically, so the edits are scripted and auditable rather than per-frame manual retouching: (i) the prompt text is standardized to the single mouse category; (ii) lead-in frames recorded before tracking has acquired every animal are dropped, since they carry missing rather than sparse labels; (iii) track ids belonging to one animal that was lost and re-acquired are merged, after checking the two ids do not overlap in time; (iv) masks are repaired by connected-component filtering and bounding box and area are recomputed from the repaired mask. Frames and identities are then reindexed from zero and the overlays reinspected.

#### Pass 2: mask-gated pose

Pose is fitted to the *curated* file rather than to the raw tracker output. Each stored segmentation is decoded back to a binary mask and the stored box and mask are passed to the gated pose head, so every keypoint is conditioned on already curated masks. Keypoints are written back in COCO person-keypoints form in flat [*x, y, v*] *→* 27 while ids, scores, track ids and masks are carried through untouched.

#### Curation 2: pose

The pass renders an annotated video in which masks, boxes and keypoints are drawn in each animal’s track colour, and every recording is reviewed on it before use. Because the raw per-keypoint confidences are retained, an over- or under-permissive visibility threshold is corrected by re-deriving *v* from the stored scores rather than by editing coordinates. Recordings whose review exposes a mask or identity fault are returned to curation 1 and processed again. Each file is finally checked against the schema, box extent and containment, polygon or RLE validity, 3 *→ K* keypoint values with *v ↓ {*0, 1, 2*}*, and agreement between `num_keypoints` and the visibility flags, before it is merged into the dataset.

### 4.5 Dataset splits

The dataset’s primary form is stored per task, isolating each recording with its own consistent image ids and frame indices, so it can be merged in different ways depending on the goal. For every evaluation in the paper the per-task recordings were merged after keeping every tenth frame, to limit near-duplicates from the 30 fps recordings.

#### Pooled

Frames from all nine assays are merged and shuffled into train, validation and test splits (70/15/15%). This split is used for the pooled MouseLite and DeepLabCut results (Figure 5a) and for the MouseLite-detection and MouseLite-segmentation variants released (Figure S5). Because frames sampled from the same recording remain only 0.33 s apart, pooled test frames closely resemble training frames; this split therefore measures in-distribution fit, and generalization is assessed with the leave-one-assay-out protocol below.

#### Leave-one-assay-out (LOAO)\

One assay is held out entirely for testing and the models are trained on the remaining assays. We evaluated two variants: *in-domain*, where validation is carved out of the eight training assays, and *out-domain*, where validation is also a held-out assay, separate from both training and test, so that seven assays are used for training. Under out-domain, the validation assay is the next one after the test assay in a fixed rotation, wrapping around, so that each assay validates exactly once across the sweep. The two gave similar results on a subsample of folds, so only out-domain LOAO is reported in the paper. Rotating the held-out assay gives nine folds, each named for the assay used as test (Figure 5). The open-field fold is excluded from the model comparison because of a software failure of the DeepLabCut cascade on this fold. Each assay is represented by a single recording, so a held-out assay is also a held-out recording, and assay-specific factors cannot be separated from recording-specific ones such as the animals, the session or the camera placement.

### 4.6 Evaluation metrics

#### 4.6.1 Cross-assignment

Cross-assignment is a label-free proxy for identity errors, defined against the segmentation rather than against keypoint ground truth, which does not exist for the UDMT clips. For each tracked instance, keypoints above confidence 0.3 are classified by the mask they fall inside: the instance’s own mask, another instance’s mask, or background. A keypoint on another animal is wrong by construction. The cross-assignment rate is pooled within each stratum, summed cross-assigned keypoints over summed valid keypoints, not averaged over detections; the on-own-mask rate is defined identically. Results are stratified by occlusion, the fraction of an instance’s box covered by other instances’ masks, in right-open bins (0–2, 2–5, 5–10, 10–20, 20–100%), and by the number of animals in the clip.

#### 4.6.2 Memory and throughput

Peak CUDA memory and wall-clock time were measured on the SAM 3 video wrapper alone: frames were decoded and passed straight to its forward pass, with no pose head, annotation or video writing. Each of the four UDMT clips was swept over sequence lengths of 200 to 1,000 frames in steps of 100, with mask-memory eviction on (64-frame window) and off, giving the two axes of Figure S1. CUDA memory was sampled once per frame and the per-run peak retained; throughput is frames divided by elapsed wall-clock. All runs used a single NVIDIA RTX 4000 Ada Generation GPU.

#### 4.6.3 Bounding box mAP

For each object category, predicted boxes were ranked by confidence and matched to ground-truth boxes using Intersection over Union (IoU). A match counted as a true positive when IoU exceeded a threshold. Precision and recall were computed across confidence levels to form a precision–recall curve, and Average Precision (AP) was taken as the area under this curve. We averaged AP over ten IoU thresholds (0.50 to 0.95, step 0.05) and all categories to obtain mAP.

#### 4.6.4 Bounding box mAR

Average Recall (AR) measured the fraction of ground-truth boxes correctly detected, given a fixed number of detections per image. AR was averaged over ten IoU thresholds (0.50 to 0.95, step 0.05) and all categories to obtain mAR@[0.50:0.95].

#### 4.6.5 Keypoint mAP

Keypoint matching used Object Keypoint Similarity (OKS) in place of IoU:

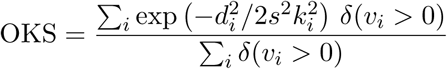

where *d_i_* is the distance between predicted and ground-truth keypoint *i*, *s* is the object scale, *k_i_* is a per-keypoint falloff constant, and *ω*(*v_i_ >* 0) is an indicator function equal to 1 when keypoint *i* is labelled and visible and 0 otherwise. We used a flat *k_i_* = 0.1 for every keypoint following (3, 12). AP was computed as for boxes and averaged over ten OKS thresholds (0.50 to 0.95, step 0.05) to obtain keypoint mAP.

#### 4.6.6 Keypoint mAR

Keypoint AR measured the fraction of ground-truth poses correctly localized, using OKS for matching. AR was averaged over the same ten OKS thresholds to obtain keypoint mAR.

### 4.7 Model training

Both models were trained with the hyperparameters listed in Table 2. All parameters not specified there were left at their respective framework defaults (RF-DETR v1.9.0 and DeepLabCut v3.0.1).

**Table 2:** Training parameters for MouseLite and DeepLabCut.

|  | MouseLite-keypoints | DeepLabCut (SuperAnimal) |
| --- | --- | --- |
| Paradigm | end-to-end prediction | top-down: Faster R-CNN + HRNet heatmaps |
| Input | $576 \times 576$ | detector $\leq 1152$ px short side; pose $256 \times 256$ animal crops |
| Epochs | 60 | 60 pose + 25 detector |
| Batch (effective) | $2 \times 4 = 8$ | 8 pose, 2 detector |
| Optimizer | AdamW, wd $1 \times 10^{-4}$, clip 0.1 | AdamW |
| Learning rate | $1 \times 10^{-4}$, cosine to 0, 1-epoch warmup | $1 \times 10^{-5}$, constant |
| Early stopping | Yes (patience 10) | No |
| Checkpoint evaluated | Lowest validation loss | Lowest validation loss |
| Seed | 0 | 42 |

### 4.8 Hardware

All experiments were run on a single NVIDIA RTX 4000 Ada Generation GPU or a single NVIDIA RTX 5070.

### 4.9 Software

Experiments were coded in Python benefiting from awesome open-source libraries: deeplabcut (1), rfdetr (31), supervision (41), opencv-python (42), transformers (43), pytorch (44), faster-coco-eval (45).

DeepLabSAM, MTMB, and MouseLite are distributed as Python packages whose dependencies are pinned in uv lock files, so that each environment is reproduced with a single command. The software was tested on Linux (Manjaro), Windows and macOS. MouseLite inference also runs without a GPU: on a laptop CPU, the joint detection–keypoint model processes an image in 603 ms (1.7 fps), and the detection and segmentation models in 105–391 ms and 155–428 ms, respectively, from nano to large.

For each video, it writes a COCO-format JSON file (boxes, 27 keypoints, detection scores, track identities and frame rate), a CSV table with one row per detection (time, track identity, centroid, box and keypoint coordinates) and a per-track summary (coverage, distance, mean and maximum speed, duration). Coordinates and speeds are expressed in pixels.

## 5 Acknowledgement

We thank the IGF animal facility and the “Physiopathology of neuronal plasticity” lab for scientific discussions.

Funding: 80PRIME “NeuroJac” from the CNRS MITI (JC, MB, JN)), Centre National de la Recherche Scientifique (CNRS UMR 5203), Université de Montpellier, Inserm (U 1191), Agence Nationale de la Recherche (ANR Timetag to JN).

## 6 Author Contribution

Conceptualization : JC, JN. Data acquisition : JC, ST. Data curation : JC, ST. Funding acquisition : MB, JN. Investigation : JC, ST. Methodology : JC. Project administration : JN. Resources : JN. Software : JC. Supervision : MB, JN. Validation : JC, MB, JN. Visualization : JC, JN. Writing — original draft : JC, JN. and Writing — review & editing : JC, ST, MB, JN.

## 7 Ethics

The study was approved by the CEEA local and national committees under the authorized establishment license B34-172-41.

## 8 Data availability

The MTMB dataset is publicly available on Hugging Face (https://huggingface.co/datasets/juancobos/MultiTaskMouseBehaviour) and archived on Zenodo (46) (https://zenodo.org/records/23035460) under a CC BY-NC 4.0 licence.

## 9 Code availability

All code developed for this study is open source and publicly available on GitHub. The code for DeepLabSAM (https://github.com/juan-cobos/DeepLabSAM) and MouseLite (https://github.com/juan-cobos/mouselite), together with the trained MouseLite weights, is released under the Apache 2.0 licence. The code used to build and curate MTMB (https://github.com/juan-cobos/mtmb) is released under the MIT licence.

---

**Figure S1:**
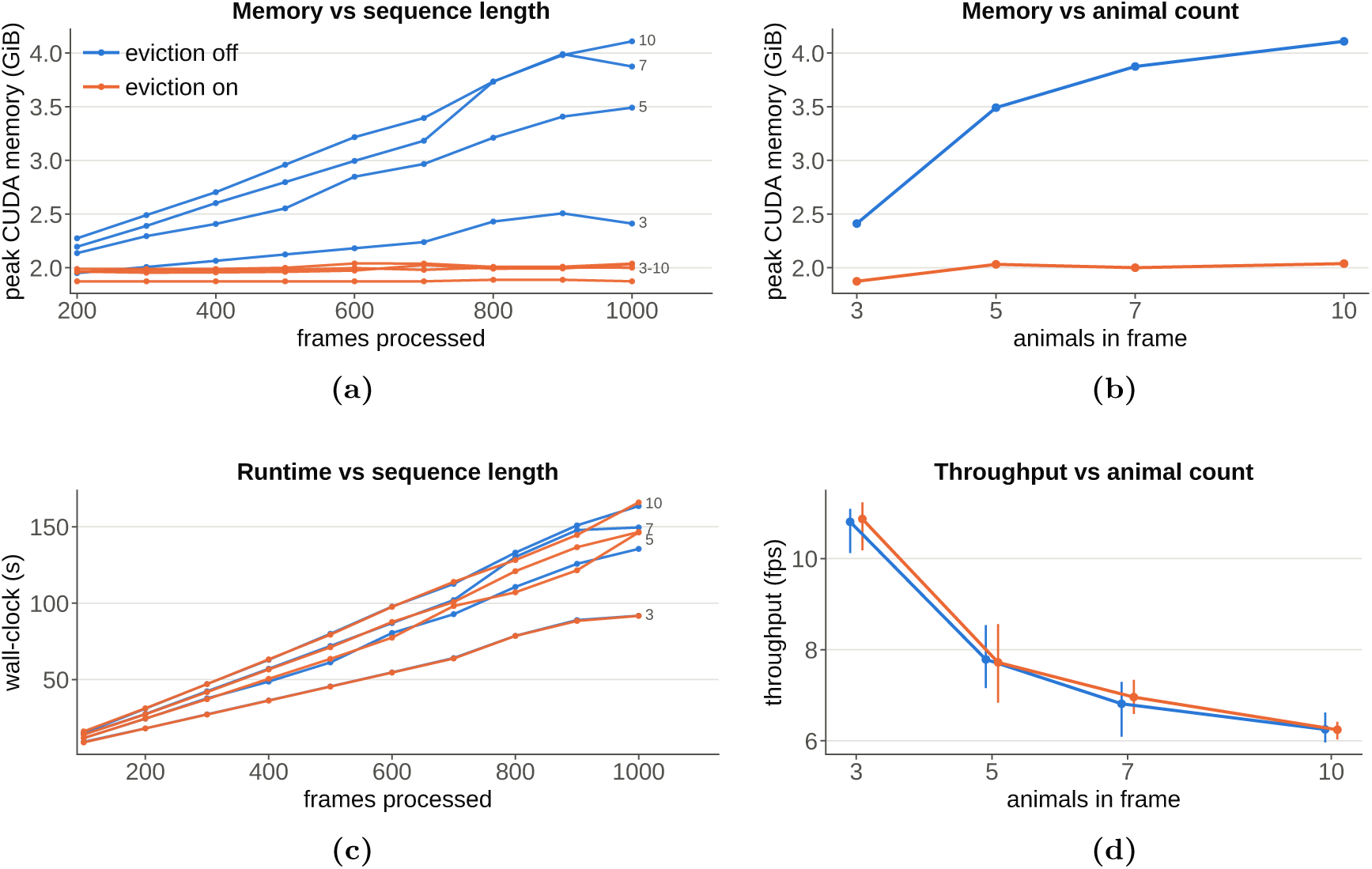
Bounded mask-memory makes long videos tractable. Peak memory and runtime measurements are performed on the four public UDMT mouse clips of 3, 5, 7 and 10 freely interacting animals (19, 20). Blue denotes mask-memory eviction disabled and orange denotes the proposed bounded mask-memory mechanism with eviction enabled. (**a**) Peak CUDA memory versus number of frames processed, one line per clip and eviction condition; the mouse count is direct-labelled at the right end of each line, with the four eviction-on curves collapsing into a single bundle labelled 3–10. With eviction disabled, memory grows linearly with sequence length; with the 64-frame mask-memory window it remains approximately constant. (**b**) Peak memory at the longest sequence (1,000 frames) as a function of the number of animals in frame: without eviction, memory increases with animal count in addition to sequence length; with eviction it remains bounded. (**c**) Wall-clock time versus frames processed, using the same encoding as in **a**. The two conditions overlap, indicating that the memory bound does not introduce measurable runtime overhead. (**d**) Throughput versus number of animals in frame, pooled over the sweep lengths (points, mean; bars, min–max across sweep lengths).

**Figure S2:**
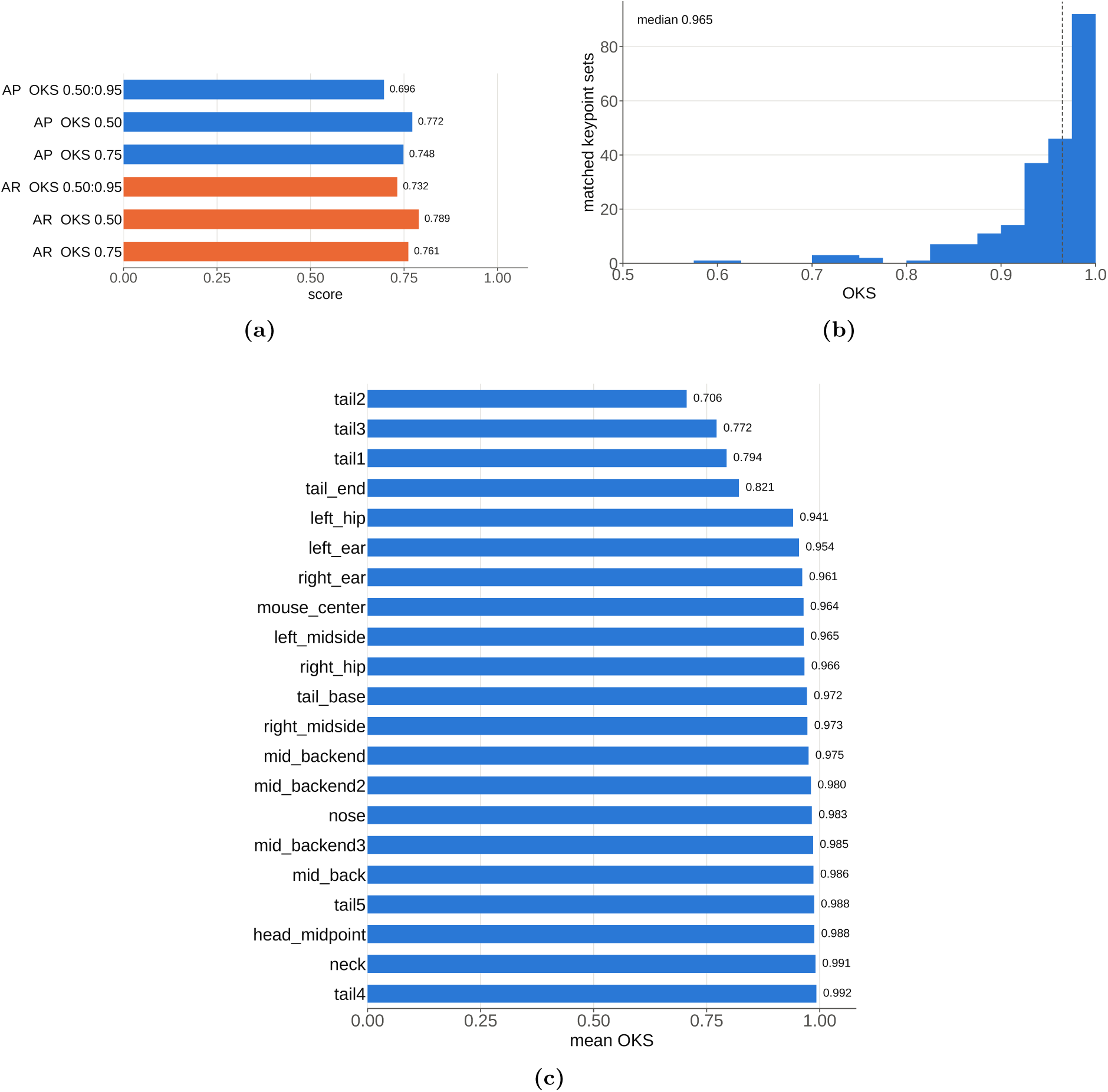
DeepLabSAM pose estimates against human-annotated ground truth. DeepLabSAM predictions scored against the human-labelled test split of SuperAnimal-TopViewMouse-5K (47). (**a**) Average precision (AP) and average recall (AR) at OKS 0.50:0.95, 0.50 and 0.75. (**b**) OKS distribution over the matched pairs (median 0.965). (**c**) Mean OKS per bodypart, worst first. Error concentrates in the tail (tail1–tail3, tail_end: 0.71–0.82), while trunk and facial landmarks are near-ceiling (*↘*0.94).

**Figure S3:**
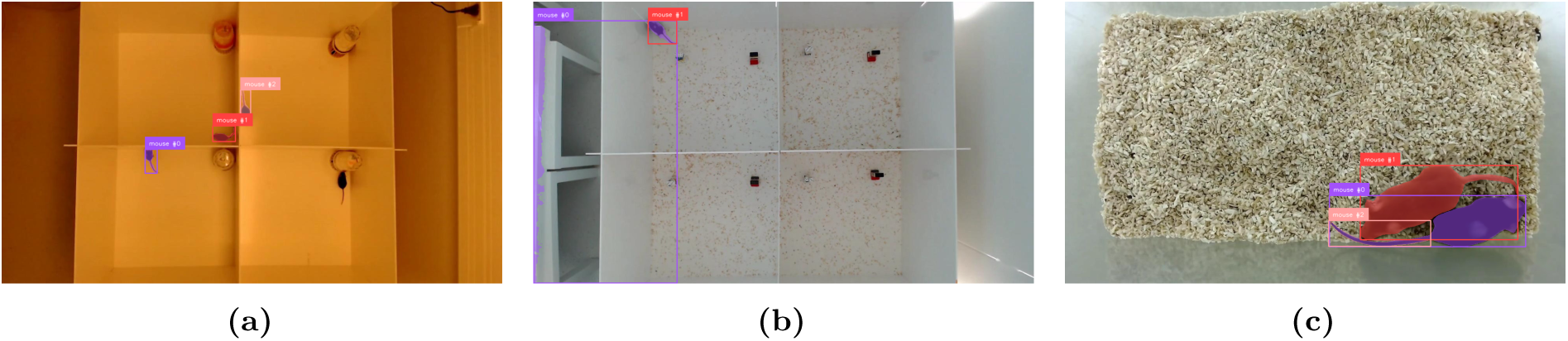
Curated examples illustrating three targeted post-processing corrections. (a) Masks from the first pass were incomplete, so the frame was re-segmented with SAM3 to recover full per-mouse silhouettes. **(b)** A single instance mask leaked across the arena into an implausibly large region; a connected-components analysis retains only the dominant component, restoring a compact mask. **(c)** Under occlusion and clutter, redundant proposals exceed the known animal count, so top-*k* selection on the segmentations keeps only the *k* highest-scoring, non-redundant instances.

**Figure S4:**
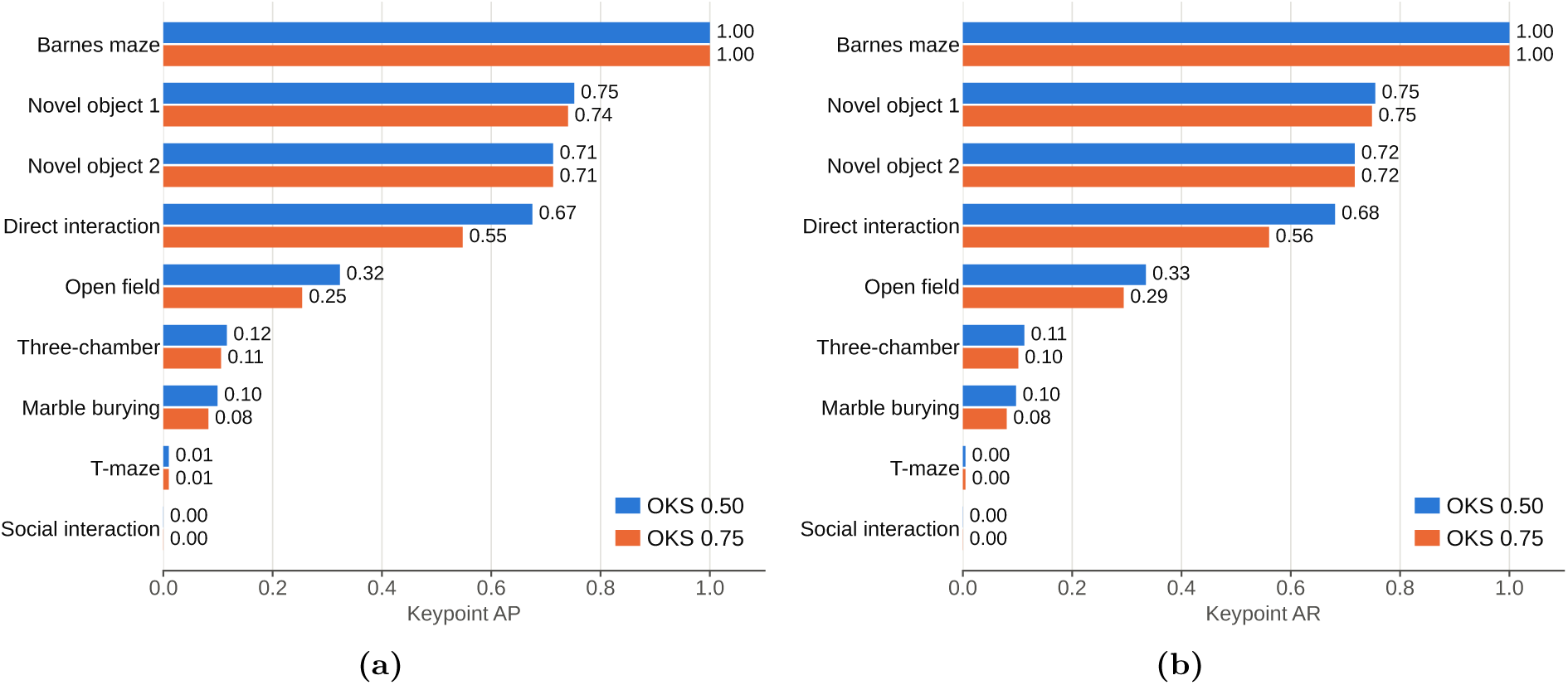
Zero-shot keypoint accuracy at permissive and strict localization thresholds. Keypoint accuracy of the pretrained DeepLabCut model on each held-out recording of the Multi-Task Mouse Behaviour dataset (Figure 4), evaluated at OKS = 0.50 (blue) and OKS = 0.75 (orange). Recordings are sorted by AP_50_. (**a**) Keypoint average precision. (**b**) Keypoint average recall. Recordings with high accuracy keep most of it at the stricter threshold (Barnes maze 1.00 at both; direct interaction 0.67 to 0.55), whereas the three-chamber test (0.12), marble burying (0.10), the T-maze (0.01) and social interaction (0.00) are already poor at OKS = 0.50. AP closely tracks AR throughout, so almost all detections that are returned are correct and the loss comes from animals that are missed, not from false positives.

**Figure S5:**
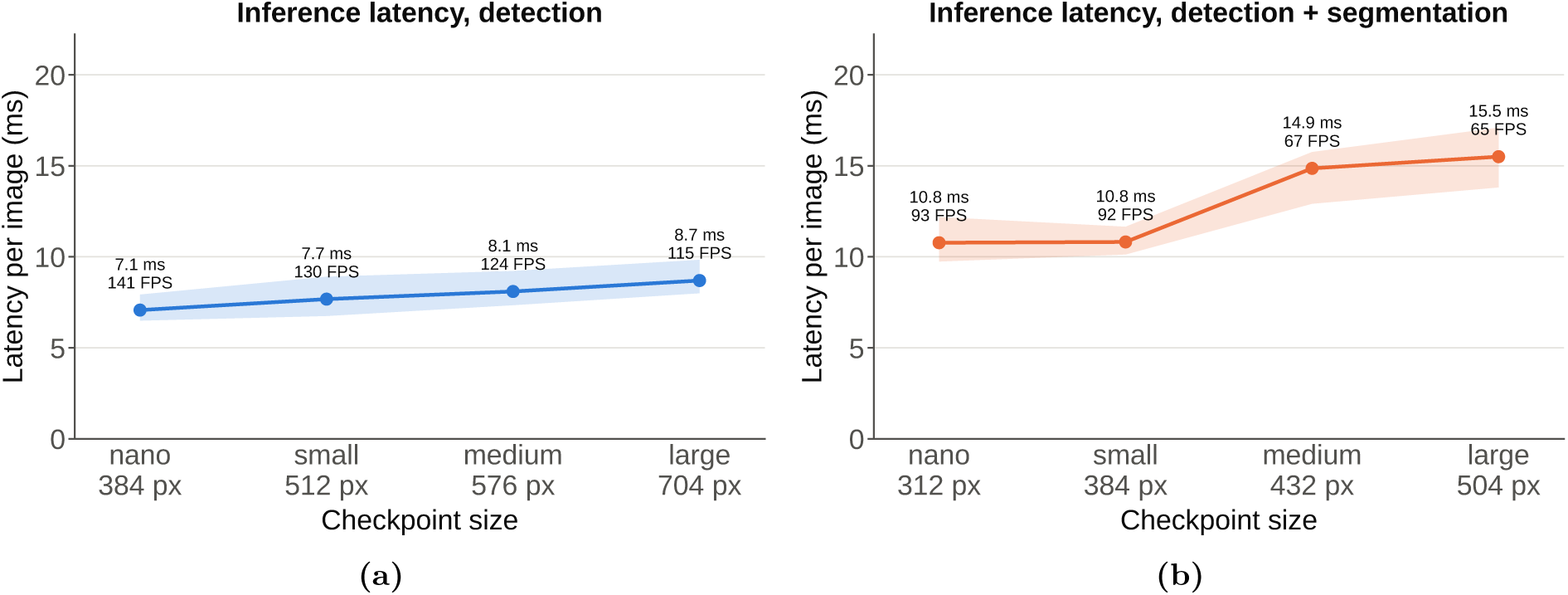
Detection and segmentation checkpoints at varying sizes. In addition to the MouseLite–keypoints model (Figure 5), we release MouseLite-detection and MouseLite–segmentation checkpoints at four scales (nano, small, medium, large), each trained on the pooled Multi-Task Mouse Behaviour dataset. Inference latency is measured per image on a single GPU; the point marks the mean and the shaded band the per-image distribution. (**a**) MouseLite–detection checkpoints, from 7.1 ms (141 FPS) at nano/384 px to 8.7 ms (115 FPS) at large/704 px. (**b**) MouseLite–segmentation checkpoints, from 10.8 ms (93 FPS) at nano/312 px to 15.5 ms (65 FPS) at large/504 px. Adding the segmentation head raises latency but every configuration still runs well above real time.

**Figure S6:**
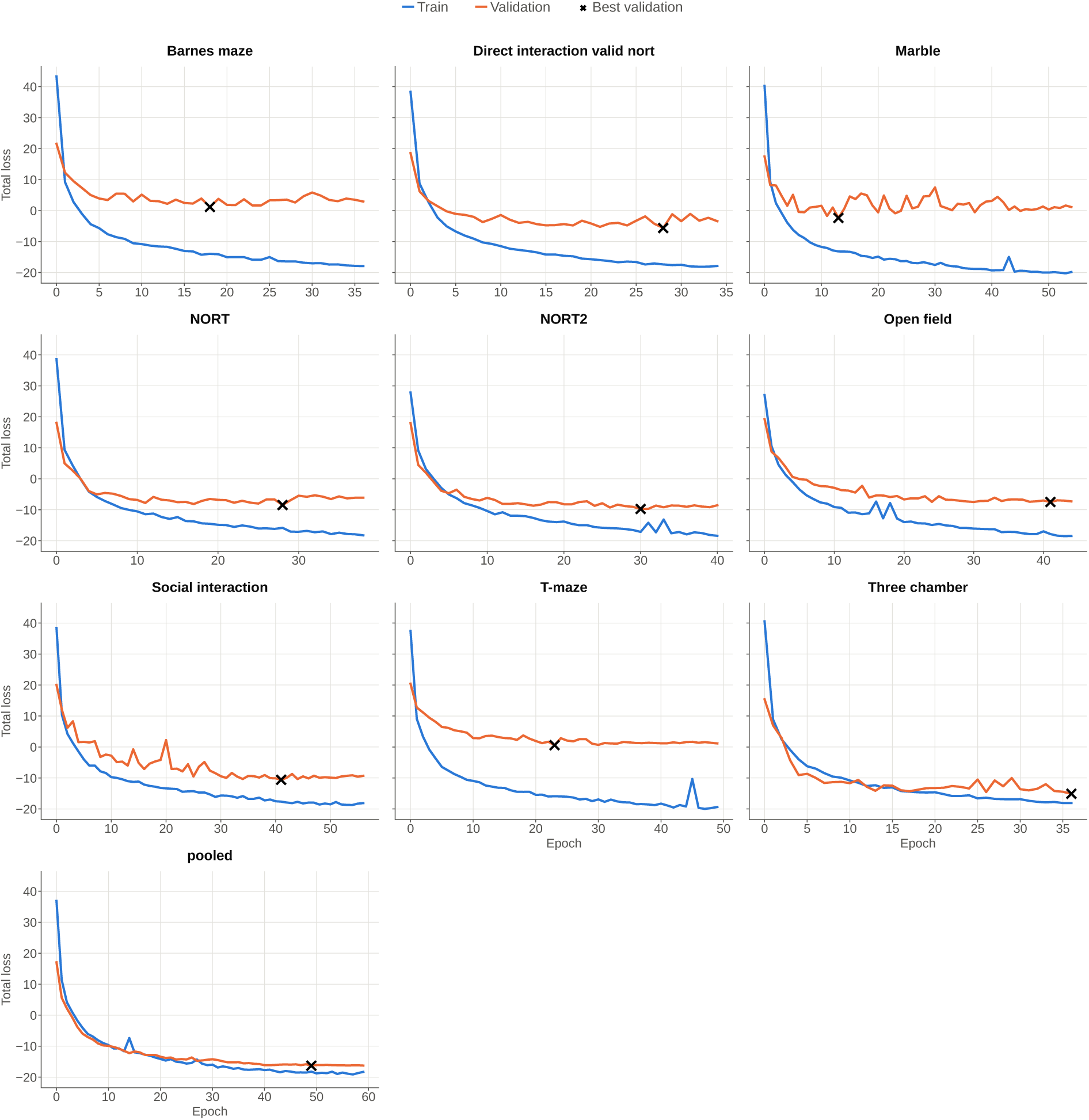
Training and validation curves for pooled and leave-one-assay-out runs (MouseLite). Total loss versus epoch, with one panel per run titled by the held-out assay, plus the *pooled* run trained on all nine recordings. Train loss is shown in blue, validation loss in orange, and the black X marks the epoch of lowest validation loss, whose checkpoint is used for evaluation.

**Figure S7:**
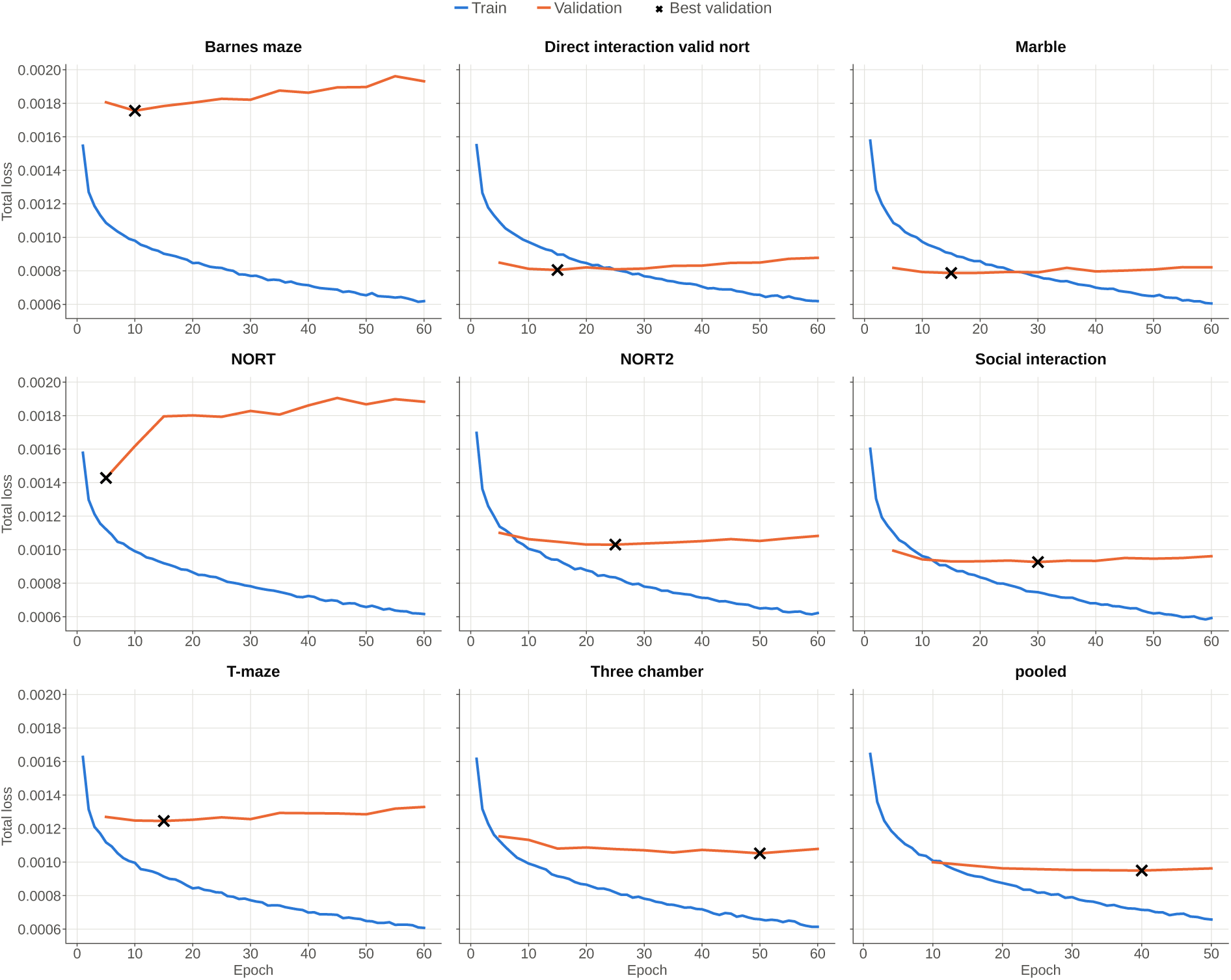
Training and validation curves for pooled and leave-one-assay-out runs (DeepLab-Cut). Total loss versus epoch, with one panel per run titled by the held-out assay, plus the *pooled* run trained on all nine recordings. Train loss is shown in blue, validation loss in orange, and the black X marks the epoch of lowest validation loss, whose checkpoint is used for evaluation. The DeepLabCut cascade (pose-head training shown). On several held-out assays (e.g. Barnes maze, NORT) validation loss rises after an early minimum while training loss keeps decreasing, a sign that fine-tuning starts fitting the eight training assays too closely at the expense of held-out generalization; the best checkpoint is therefore reached early.

